# Campylobacter jejuni Infection Is Associated With Cell-Cycle, Redox, and Metabolic Remodeling of Human Intestinal Epithelial Cells at Single-Cell Resolution

**DOI:** 10.64898/2026.09.04.749441

**Authors:** Angela Gao, Violet Newhart, Selena Kassis, Paloma Bravo, Ashfaqul Alam

## Abstract

*Campylobacter jejuni* is the leading cause of bacterial enteric infections worldwide, including in the US. It is also a zoonotic pathogen that is transmitted by food and water, causing diarrhea and intestinal inflammation, and is responsible for high morbidity and mortality in young children, the elderly, and immunocompromised patients. During infection, *C. jejuni* profoundly perturbs intestinal epithelial physiology, leading to widespread mucosal damage and inflammatory response. However, the heterogeneity and coordination of host transcriptional responses remain incompletely defined. We performed single-cell RNA sequencing of Caco-2 intestinal epithelial cells from uninfected Control and *C. jejuni*-infected biological samples using 10x Genomics GEM-X Flex chemistry. Our dataset comprised 159,088 cells and 18,142 measured features. A Seurat workflow resolved 19 transcriptional states and revealed a highly reproducible condition-associated redistribution of clusters across biological replicates. Our study demonstrated that C. jejuni infection was associated with increased representation of G2/M-classified cells and coordinated induction of checkpoint and mitotic-spindle genes, including CDKN1A, WEE1, MAD2L1, BUB1B, PLK1, CDC20, CDK1, and UBE2C. Furthermore, ranked enrichment independently identified a C. jejuni-enriched Reactome mitotic spindle checkpoint program (NES ∼1.67, FDR ∼0.013), while Hallmark analysis identified strong TNF/NF-kB, hypoxia, G2M checkpoint, TGF-beta, glycolysis, apoptosis, p53, and mTORC1-associated programs. Infection was also associated with significantly decreased JDP2 and higher NOX1 and DUOX2 gene expression. Furthermore, scMetabolism/AUCell identified 20 KEGG metabolic pathways at FDR <0.05 in a balanced 10,000-cell analysis, with 12 lower and 8 higher in infection. Together, these data support a model in which *C. jejuni* dramatically restructures epithelial transcriptional states and is accompanied by mitotic-checkpoint, inflammatory/stress, redox, and metabolic remodeling.

## Introduction

*Campylobacter jejuni* is a principal cause of bacterial gastroenteritis, with 400 million cases worldwide and 1.5 million in the U.S. annually^1,2^. Infection can also lead to late-onset diseases such as Guillain-Barré Syndrome, colorectal cancer, inflammatory bowel disease, irritable bowel syndrome, and celiac disease^1,3^. According to the CDC, these infections cost up to $6.8 billion each year in the U.S. Human infection often occurs through the ingestion of contaminated food, especially poultry products, which leads the bacterium to colonize the colon and cause severe inflammatory and bloody diarrhea, high fever, vomiting, severe abdominal pain, and severe mucosal damage. In some patients, especially immunocompromised individuals, the elderly, and children, infection may progress to bacteremia or systemic infection and, in some cases, may cause mortality. Importantly, *C. jejuni* has been designated a serious antibiotic resistance threat by both WHO & CDC. Despite the high prevalence of human campylobacteriosis in the USA & worldwide, and the emergence of antibiotic resistance in clinical isolates, the mechanisms by which *C. jejuni* regulates its pathogenesis in the host and disease severity remain incompletely understood.

Several virulence factors of *C. jejuni* have been identified that contribute to colonization, host invasion, immune evasion, and disease pathogenesis^4–5^. These include adhesion proteins such as CadF and FlpA, the cytolethal distending toxin (CDT), invasion-associated proteins including Cia proteins, flagella-dependent motility and secretion, chemotaxis, oxidative stress defense system (KatA, SodB, AphC), iron acquisition systems, multidrug resistance system, and metabolic pathways required for adaptation to the intestinal environment. Despite extensive research, major gaps remain in understanding how the human gut microbiota influences *C. jejuni*’s response to the host and regulates expression of virulence factors and infection dynamics^7–11^.

A recent scRNA sequencing study suggested that *C. jejuni* infection alters host cell-cycle progression and is associated with transcriptional programs involving PLK1 signaling, mitosis, hypoxia/HIF signaling, inflammatory pathways, p53-associated responses, and focal adhesion. This study provides an important external framework for evaluating whether related programs are evident in an independent epithelial model.

In the present study, we analyzed six Caco-2 epithelial samples comprising three uninfected controls and three *C. jejuni*-infected samples in a single 10x Genomics Flex experiment. We aimed to define infection-associated transcriptional states across the whole epithelial population, determine whether altered cell-cycle composition accompanies infection, identify transcriptional differences within phase-matched populations, and discover and validate infection-responsive genes and pathways. We additionally prioritized JDP2, NOX1, and DUOX2 because independent experimental observations highlight a potential regulatory relationship between JDP2 and epithelial oxidase expression during *C. jejuni* invasion of intestinal epithelial cells.

## Materials and Methods

### Cell culture and experimental design

Caco-2 intestinal epithelial cells were cultured in MEM containing 10% fetal bovine serum overnight. Cultures were rinsed/released into MEM containing 1% fetal bovine serum and either maintained as uninfected Controls or infected with *C. jejuni* for 24 h. Cells were passed through a 30-micrometer strainer and enumerated with an EVE cell counter before downstream processing.

### Single-cell library preparation and sequencing analysis

Libraries were generated for six samples using Chromium GEM-X Flex Fixed RNA Profiling - Human, v2 chemistry with the Chromium Human Transcriptome Probe Set v2, GRCh38-2024-A. We processed all samples in a pooled Flex workflow and demultiplexed them using sample-specific probe barcodes.

The integrated feature-barcode matrix contained 18,132 genes and 159,088 cells. Barcode-level metadata from the 10x graph-based clustering export and condition assignment file matched the matrix exactly, with no duplicated barcodes. Cell Ranger multi identified 26,035, 27,975, and 23,524 cells for the three uninfected control samples, respectively, and 23,066, 31,022, and 27,466 cells for the three C. jejuni-infected samples, respectively, yielding 159,088 cells across the six samples included in the analysis. Condition totals were 77,534 uninfected and 81,554 C. jejuni-infected cells.

### Seurat preprocessing, dimensional reduction, and clustering

Analyses were performed in R 4.3 using Seurat 5.3. RNA counts were normalized with NormalizeData (scale factor 10,000), 2,000 variable features were selected, and PCA was performed. The first 18 principal components were used for neighbor graph construction, UMAP, and clustering. Harmony was deliberately not applied so that condition-associated structure was not removed. Clustering at resolution 1.1 generated 19 transcriptional states.

### Cluster markers and quality assessment

Positive markers for all 19 clusters were identified using Seurat FindAllMarkers with min.pct = 0.10, log2 fold-change threshold = 0.25, and Wilcoxon testing. The marker table contained 23,977 rows. Cluster-level nCount_RNA and nFeature_RNA distributions were examined. We calculated mitochondrial-probe signal using 12 detected mitochondrial genes.

### Cell-cycle scoring

Cell-cycle state was inferred with Seurat S- and G2/M-phase gene sets; 42/43 S-phase genes and all 54 G2/M genes were represented. The current scored object contained 45,877 G1, 55,863 G2/M, and 57,348 S-classified cells. We summarized condition-, cluster-, and replicate-level phase distributions.

### Condition-level differential expression

Seurat FindMarkers with the Wilcoxon test was used with logfc.threshold=0, and min.pct=0, testing all 18,132 genes. Positive avg_log2FC indicates higher expression in *C. jejuni*. BH-adjusted P values were calculated. We summarized BecDiscovery lists at 1.4-fold thresholds.

### Pathway enrichment and sample-level pathway visualization

Direction-specific differential-expression lists were analyzed with enrichR/BioPlanet_2019 for comparison with Talukdar et al. Cutoff-free ranked GSEA was additionally performed with fgsea against human Reactome and MSigDB Hallmark gene sets using all 18,132 genes ranked by *C. jejuni*-versus-Control avg_log2FC. Positive NES indicates enrichment toward C. jejuni. Leading-edge detection frequencies were examined to identify sparse-gene artifacts. Metabolic pathway activity was quantified with scMetabolism KEGG gene sets and AUCell in a balanced 10,000-cell subset (∼1,666-1,667 cells/sample). AUCell scores were averaged by biological sample and compared between three Control and three C. jejuni samples using Welch t tests with BH correction across 85 pathways.

## Results

### A large epithelial single-cell dataset captures condition-associated transcriptome reorganization at the single-cell level

One profound limitation of bulk RNA sequencing is that it averages physiological changes across a population of cells. Hence, to dissect the direct interactions of *C. jejuni* with intestinal epithelial cells that may progress through stages; to investigate the heterogeneity of transcriptional states; to determine the pre-existing or infection-associated states enriched among responsive cells; to identify transcriptional states to study physiological and metabolic changes; and to explore the infection outcomes, we have infected the Caco-2 cells with *C. jejuni* for 24 hours. We processed all samples within a pooled 10X Genomics Flex workflow and demultiplexed them using sample-specific probe barcodes. Libraries were generated using Chromium GEM-X Flex Fixed RNA Profiling - Human, v2 chemistry with the Chromium Human Transcriptome Probe Set v2.0.0, GRCh38-2024-A. The final dataset comprised 159,000 8 Caco-2 epithelial cells from six biological samples.

Sequencing was performed on the Illumina NovaSeq Platform, 25B flow cells, 2.9 billion total reads, and 159,000 cells with about 9,000 sequence reads per cell. Reads mapped to about 97% of the probe set, with 93.7% of reads mapped to cells. We pooled barcoded libraries from 6 samples into one gene-expression library. We performed the analysis in an R-based workflow using the Seurat platform. RNA counts were normalized with NormalizeData (scale factor 10,000), 2,000 variable features were selected, and PCA was performed (Figure 1). The PCA plot showed a pronounced *C. jejuni* infection-associated organization in the global transcriptional state in Caco-2 cells. As shown in Figure 1A, the two conditions form a strong diagonal gradient across the 1^st^ two principal components. However, the populations are not completely separated; rather, a substantial region of overlap remains in the center, with substantial heterogeneity and retention of shared cellular states. Thus, we predict that *C. jejuni* triggered a broad and detectable reorganization of the cellular transcriptional landscape. We used the first 18 principal components to construct the neighbor graph and run UMAP analysis (Figure 1B). Similarly, the UMAP analysis revealed condition-associated organization of the Caco-2 transcriptional landscape. Furthermore, shared-nearest-neighbor clustering identified 19 transcriptional clusters. Replicate-level composition was highly reproducible: Control-associated clusters included 0, 5, 7, 9, 10, 12, and 13, whereas C. jejuni-associated clusters included 1, 2, 4, 6, 8, and 11. Interestingly, cluster 0 represented about 23.8-25.0% of each Control sample but only about 0.3-0.7% of infected samples. Cluster 7 represented about 11.9-12.5% of Controls but about 0.1-0.2% of infected samples. Conversely, clusters 1, 2, 4, 6, and 8 expanded reproducibly in infection, demonstrating broad condition-associated restructuring of epithelial transcriptional states. Thus, this analysis allowed us to determine whether *C. jejuni* produces a uniform epithelial response or generates distinct susceptible, inflammatory, metabolically altered, barrier-disrupted, and surviving cellular states and to identify the molecular and transcriptional programs associated with each outcome.

**Figure 1.**
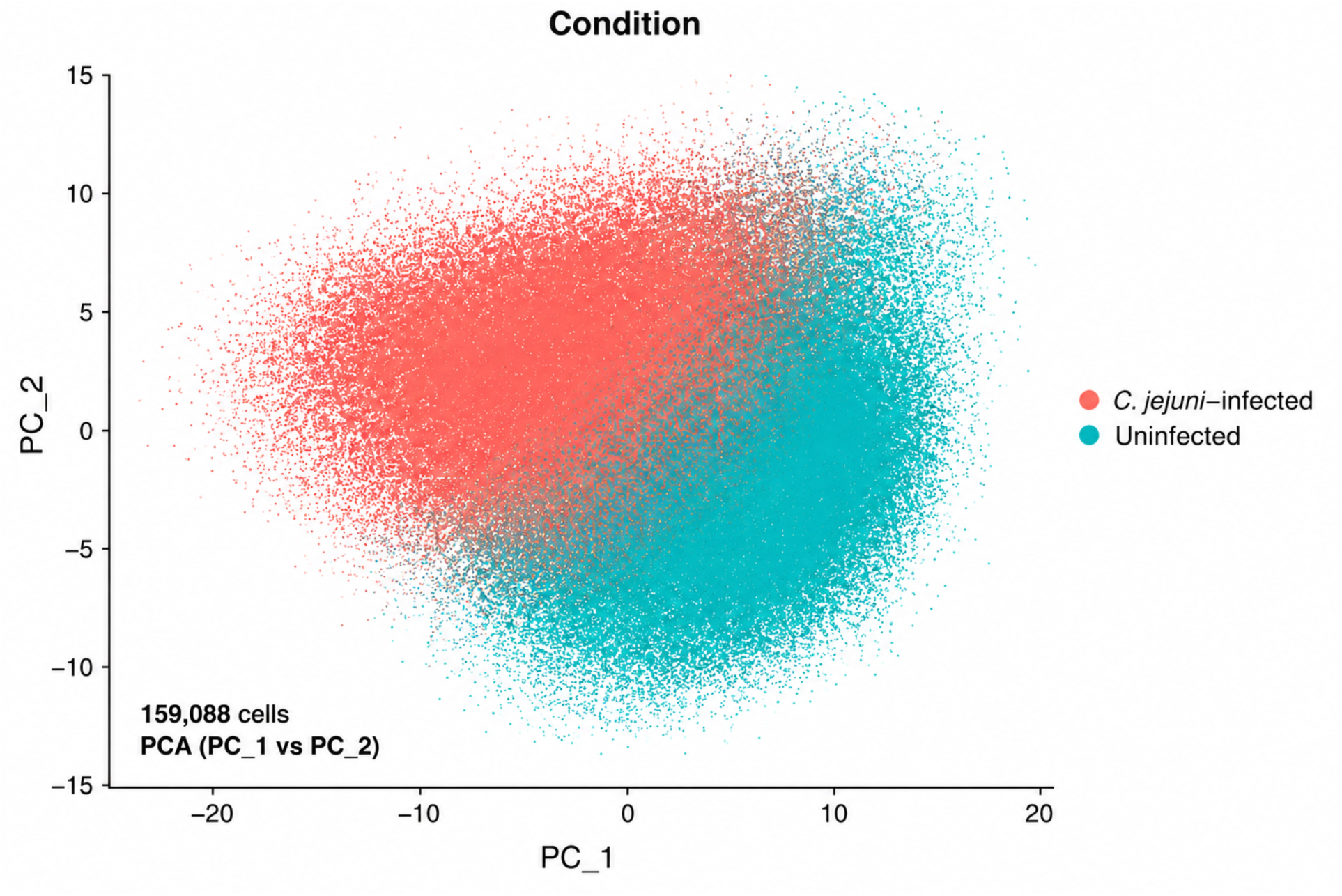
Global structure and replicate consistency of the Seurat analysis. PCA and UMAP visualize Caco-2 cells by infection condition, the independently generated 19-cluster solution (resolution 1.1). The cluster-composition heatmap summarizes condition/sample enrichment across the 19 transcriptional states. Condition-associated structure is reproducible across biological replicates. Clusters are interpreted as transcriptional states rather than distinct cell types or directly invaded versus bystander cells. These panels are from the current Seurat analysis; Loupe Browser figures are not used.

**Figure 2.**
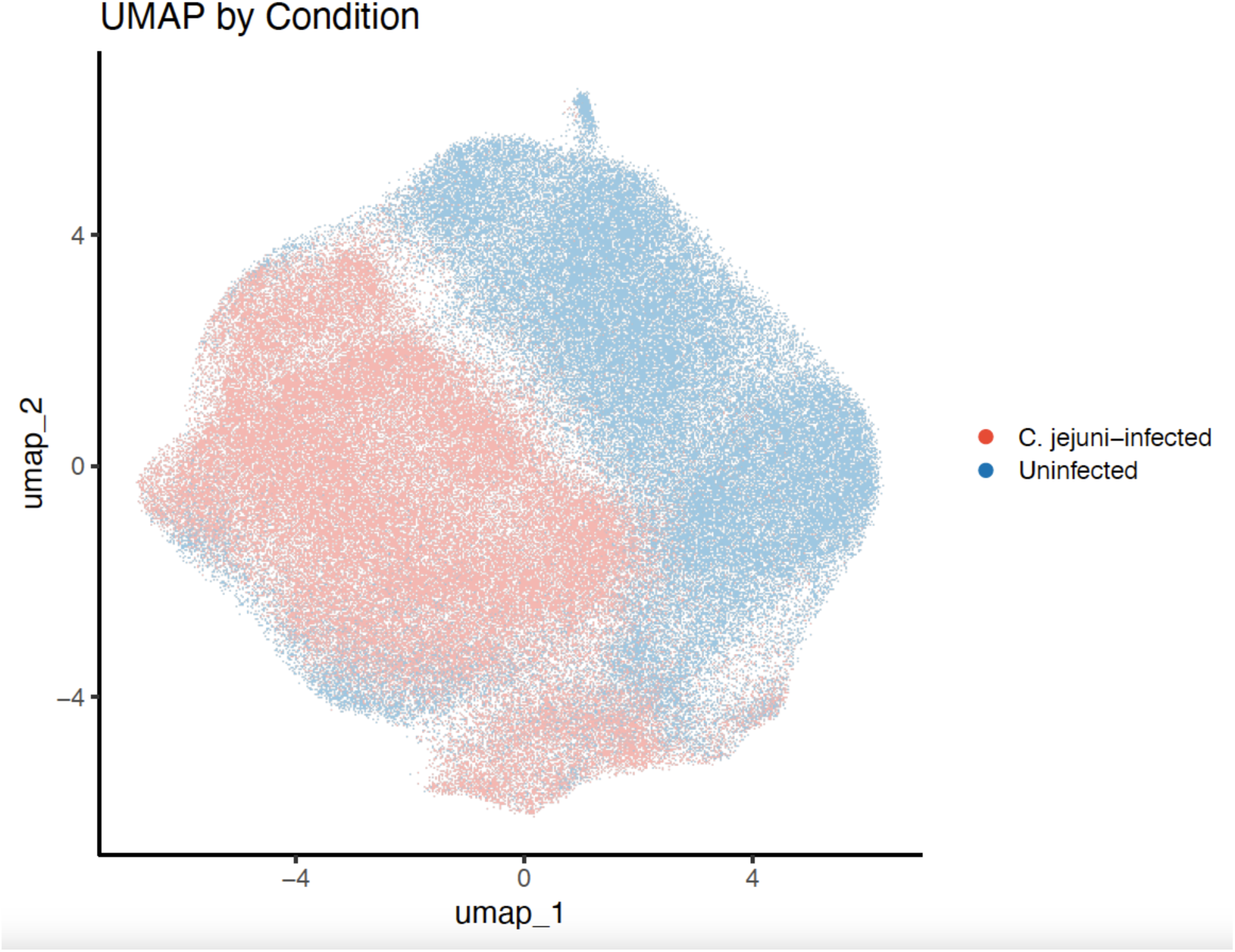
Cell-cycle remodeling associated with C. jejuni infection. Cell-cycle remodeling associated with C. jejuni infection. Cell-cycle phase was inferred using Seurat gene sets (42/43 S-phase genes and 54/54 G2/M genes represented). Current scoring classified 45,877 cells as G1, 55,863 as G2/M, and 57,348 as S. Condition-level and replicate-level phase composition, UMAP by phase, phase composition across clusters, and phase-matched Control/CJ cluster pairs. Phase is transcriptionally inferred; enrichment of a G2/M-associated program is not interpreted as proof of G2/M arrest.

**Figure 3.**
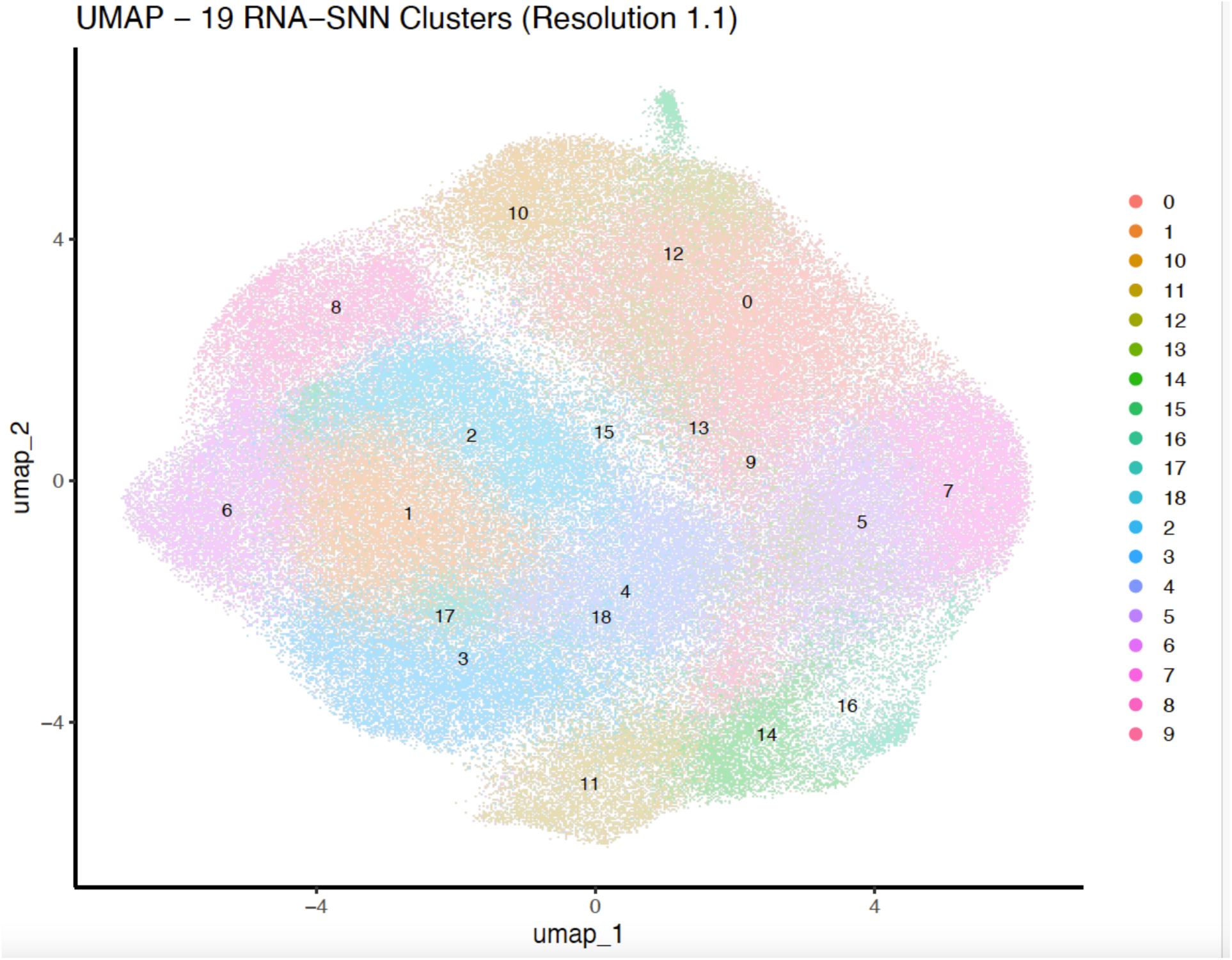
Transcriptional markers of the 19 epithelial states. Marker programs of the 19 independently defined epithelial transcriptional states. Positive markers were identified with FindAllMarkers (min.pct = 0.10; log2FC threshold = 0.25; Wilcoxon), yielding 23,977 marker rows. A top-marker heatmap, selected marker DotPlot, and JDP2/NOX1/DUOX2 expression across clusters, conditions, and biological samples.

**Figure 4.**
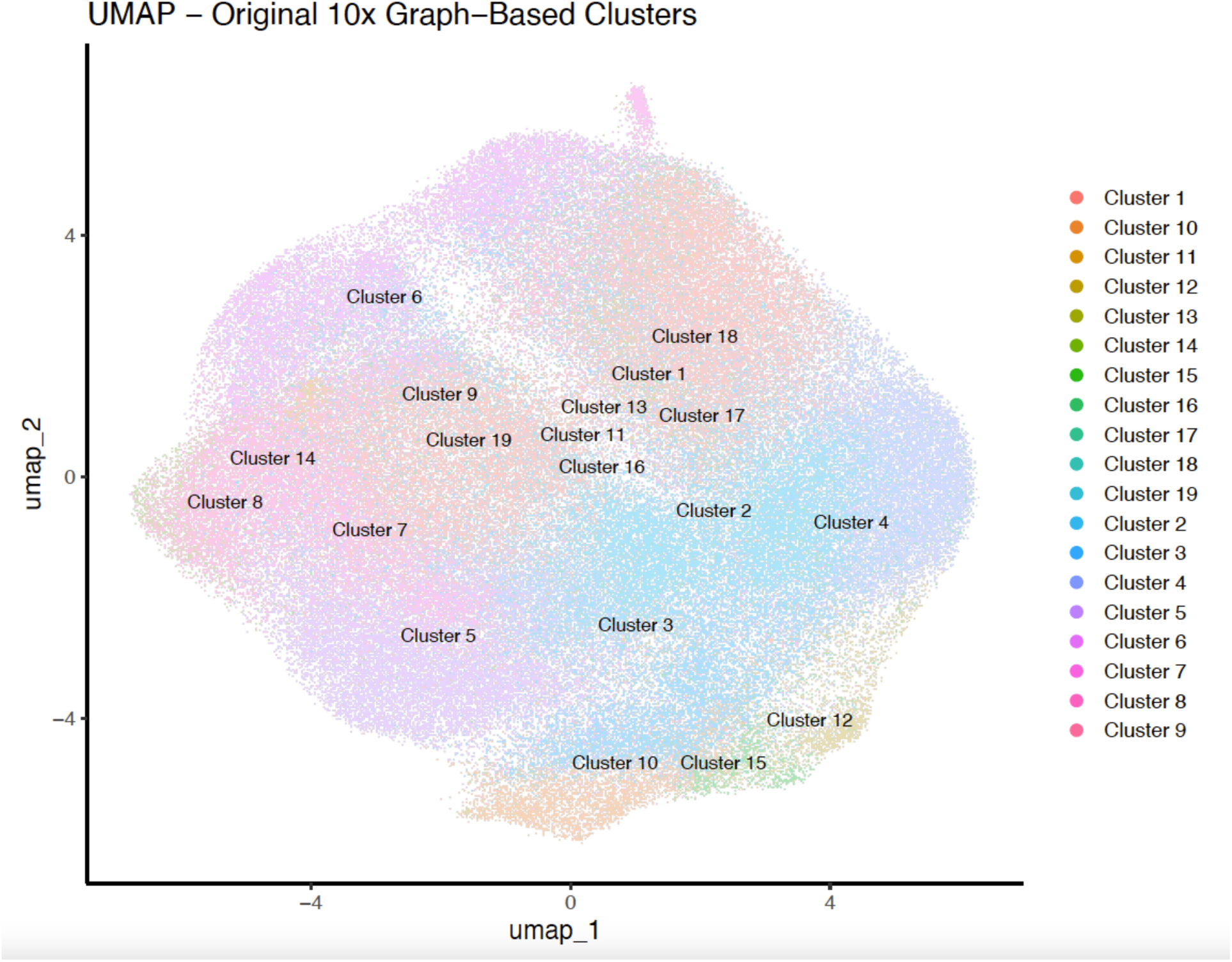
Genome-wide differential expression between C. jejuni and Control. Genome-wide single-cell differential-expression analysis comparing 81,554 *C. jejuni*-infected with 77,534 uninfected cells. Seurat FindMarkers with Wilcoxon testing, logfc.threshold = 0, and min.pct = 0.

**Figure 5.**
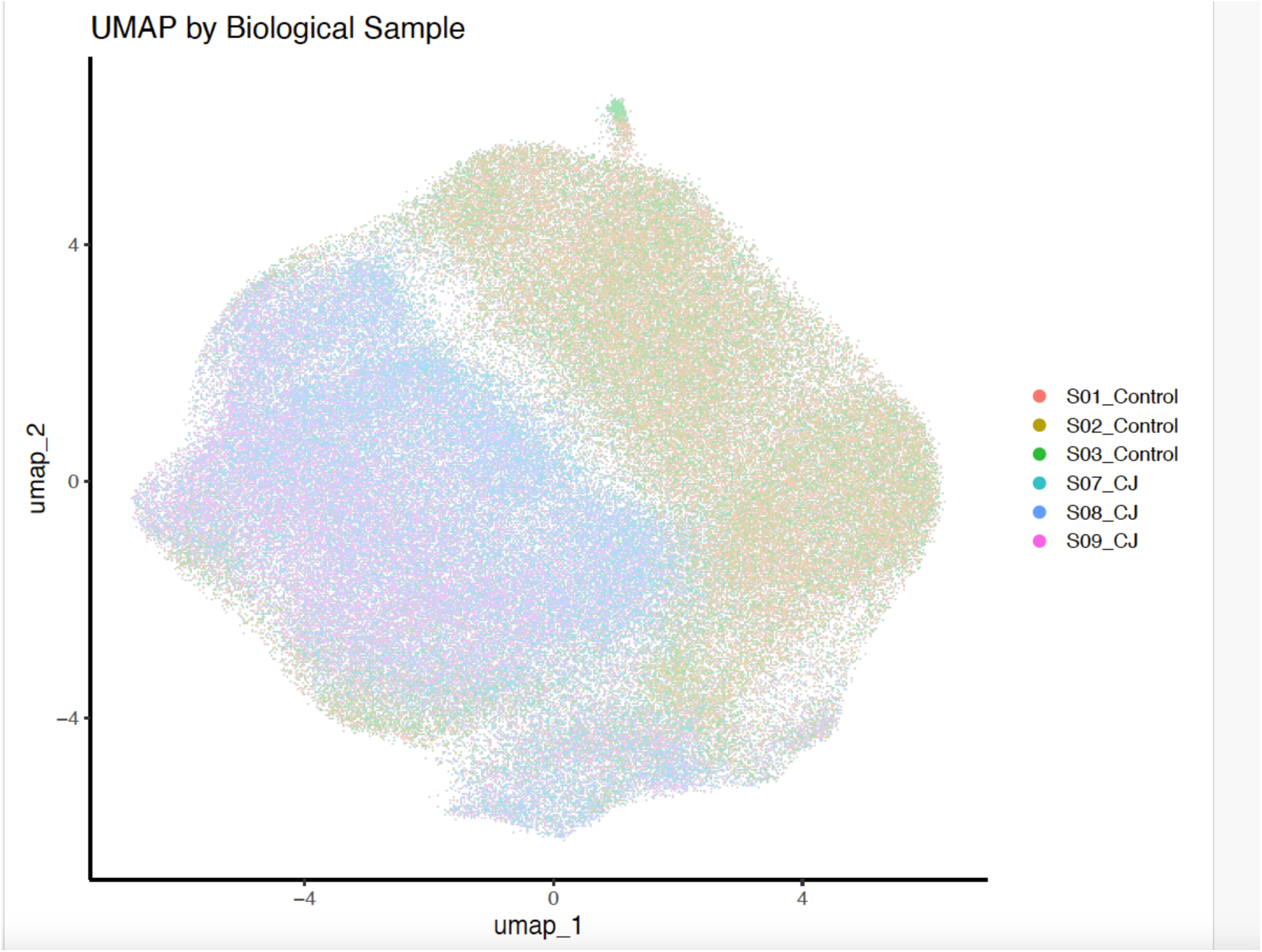
Pathway programs associated with infection. Pathway programs associated with *C. jejuni* infection. CJ-up and Control-up gene sets analyzed using BioPlanet_2019 through Enrichr/enrichR, with complementary pathway resources. Selected pathway genes were aggregated by biological sample and visualized as standardized sample-level heatmaps.

**Figure 6.**
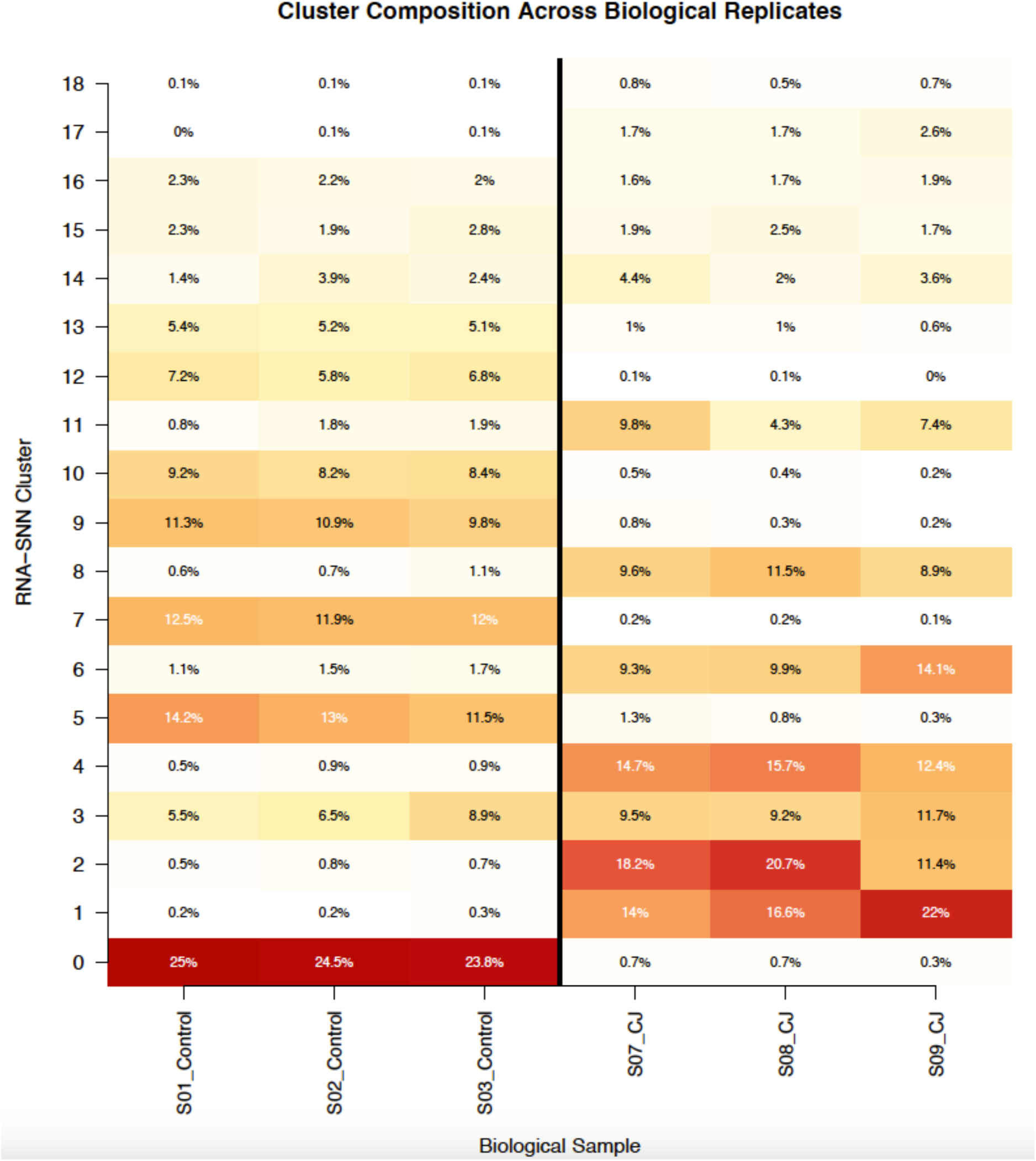
Phase-matched infection-associated remodeling. Phase-matched infection-associated remodeling. The focused G2/M comparison contrasts CJ-associated cluster 8 with Control-associated cluster 10. Adjusted-significant differences include TP53 and BUB1B (higher in cluster 8), PLK1 (higher in cluster 8), and CDC25B and GADD45A (lower in cluster 8).

Whole-dataset single-cell differential expression tested all 18,132 genes. At the 1.4-fold discovery threshold, 2,160 genes were retained (1,163 higher and 997 lower in C. jejuni); at 1.5-fold, 1,613 genes were retained; and at 2-fold, 595 genes were retained (312 C. jejuni-up and 283 Control-up). Priority genes included CDKN1A (+1.354 log2FC), DUOX2 (+1.055), NOX1 (+1.015), and JDP2 (-0.998).

Ranked Reactome GSEA identified distinct pathway-level transcriptional responses to Campylobacter jejuni infection in Caco-2 cells. Seven pathways were significantly enriched in infected cells, including RAF-independent MAPK1/3 activation, NGF-stimulated transcription, nuclear kinase/transcription-factor activation, TGF-β receptor signaling, mitotic spindle checkpoint, basigin interactions, and extracellular matrix organization. Among these, RAF-independent MAPK1/3 activation showed the strongest positive enrichment (NES = 2.32), while the mitotic spindle checkpoint was significantly enriched (NES = 1.67, FDR = 0.013) and contained 53 leading-edge genes. Together, these findings indicate that *C. jejuni* infection induces coordinated changes involving MAPK/transcriptional signaling, cell-cycle checkpoint regulation, TGF-β signaling, and extracellular-matrix remodeling.

### Pathway analysis defines major host programs associated with infection

BioPlanet analysis of the 1.4-fold lists identified 79 significant C. jejuni-associated and 7 Control-associated pathways at FDR <0.05. C. jejuni-associated programs included HIF-1/HIF-2, p53, focal adhesion, ECM-receptor, cytokine-receptor, interleukin/apoptosis, AP-1, TGF-beta, EGFR, and RAGE-related signaling. Furthermore, we found that Ranked Reactome GSEA identified seven robust C. jejuni-enriched pathways, including RAF-independent MAPK1/3 activation, nuclear kinase and transcription factor activation, TGF-beta receptor signaling, ECM organization, and the Mitotic Spindle Checkpoint (NES ∼1.67; FDR ∼0.013). Finally, the Hallmark GSEA identified 22 significant pathways, all *C. jejuni*-enriched, including TNFA/NF-kB, hypoxia, G2M checkpoint, EMT, TGF-beta, glycolysis, apoptosis, E2F targets, p53, unfolded-protein response, mitotic spindle, mTORC1, inflammatory response, and apical junction.

### *C. jejuni* infection is associated with redistribution of cell-cycle transcriptional states

Cell-cycle scoring revealed pronounced condition-associated differences. Earlier condition summaries showed ∼40.9% G2/M-classified cells in C. jejuni versus ∼28.9% in Controls, with lower G1 (∼24.9% vs ∼32.3%) and modestly lower S (∼34.2% vs ∼38.8%). The G2/M increase was reproducible across all three infected samples (∼39.7-41.7%) compared with Controls (∼28.4-29.5%). These data support accumulation of a G2/M-associated transcriptional program, indicating potential cell-cycle arrest.

Control-associated cluster 0 and CJ-associated cluster 1 were predominantly S phase; Control-associated cluster 10 and CJ-associated cluster 8 were strongly G2/M; and Control-associated cluster 5 and CJ-associated cluster 4 were predominantly G1. A focused comparison of G2/M clusters 8 versus 10 identified adjusted-significant differences in TP53, BUB1B, PLK1, CDC25B, and GADD45A. These data indicate that the pattern was not a simple canonical DNA-damage/G2-arrest signature and therefore supports a distinct infection-associated G2/M transcriptional state without establishing toxin-mediated arrest.

### JDP2, NOX1, and DUOX2 define a reproducible infection-associated expression axis

Whole-population differential expression showed JDP2 lower in *C. jejuni* (log2FC ∼-1.00; ∼0.50-fold), while NOX1 (∼+1.01; ∼2.02-fold) and DUOX2 (∼+1.05; ∼2.08-fold) were higher. These directions persisted across all three biological replicates and inferred cell-cycle phases. A broader AP-1/immediate-early panel showed JDP2 decreasing while FOSL1, ATF3, JUNB, FOSB, and other AP-1-associated genes increased, supporting coordinated stress-network remodeling without establishing JDP2 causality.

## Discussion

This study reveals extensive *C. jejuni*-associated remodeling of the Caco-2 epithelial transcriptome. The infection effect is evident in dimensional reduction, cluster composition, cell-cycle state, genome-wide differential expression, ranked pathway analysis, and metabolic pathway scoring. The strongest evidence comes from concordance across analytical levels and reproducibility across the three biological replicates.

Our data suggest that cell-cycle remodeling is supported by altered phase composition, increased expression of checkpoint/mitotic genes, and significant ranked enrichment of the Reactome Mitotic Spindle Checkpoint program. CDKN1A (∼2.56-fold), MAD2L1 (∼1.73-fold), WEE1 (∼1.72-fold), UBE2C, BUB1B, PLK1, NDC80, CDC20, and CDK1 were higher in *C. jejuni*. Together, these findings support *C. jejuni*-associated checkpoint/kinetochore remodeling and accumulation of G2/M-classified cells rather than a generalized increase in proliferation.

Finally, our data demonstrated G2/M-associated states. *C. jejuni* CDT, particularly CdtB, is known to produce DNA-damage-associated G2/G2-M blockade, including in Caco-2 models. However, our experiment used intact bacteria and did not include a cdt mutant or purified toxin. The transcriptomic phenotype is therefore compatible with known *C. jejuni* cell-cycle perturbation but cannot be attributed specifically to CDT.

Furthermore, the JDP2-NOX1-DUOX2 axis provided a second mechanistic lead. Interestingly, we found that C. jejuni infection is associated with lower JDP2 and higher NOX1 and DUOX2 across samples and phases. Independent experimental observations linking lower JDP2 with increased NOX1/DUOX2 provide a rationale for future testing; further studies are needed to demonstrate that JDP2 directly mediates the infection response. The concurrent AP-1/immediate-early signature supports broader stress-responsive transcriptional remodeling.

Metabolic analysis indicates bidirectional rather than global metabolic suppression. Twenty of 85 KEGG pathways were significant by replicate-level AUCell analysis. We have found that genes involved in glutathione metabolism were ∼14.8% lower in *C. jejuni*-infected cells (FDR ∼0.046), and 32 of 48 measured pathway genes were lower, although GCLC, GPX2, GGT5, GSTM3, and other genes increased. Furthermore, our analysis revealed that lower pentose-phosphate activity and increased NOX1/DUOX2 together generate a testable redox-homeostasis hypothesis.

Finally, we identified several limitations of the current study, including the use of a transformed epithelial cell line, transcriptionally inferred rather than flow-cytometrically measured cell-cycle phase, the absence of direct CDT perturbation, and pathway scores that do not measure metabolic flux.

Measurement of ROS, GSH/GSSG, NADPH, or metabolic flux will further strengthen these findings.

## Conclusions

Taken together, our study demonstrated that *C. jejuni* infection is associated with reproducible remodeling of Caco-2 epithelial transcriptional states, including increased representation of G2/M-classified cells, coordinated induction of checkpoint/mitotic genes, significant mitotic spindle-checkpoint enrichment, broad inflammatory/hypoxic/stress signaling, lower JDP2 with higher NOX1 and DUOX2, and bidirectional metabolic remodeling that includes altered glutathione and pentose-phosphate programs. These findings define a *C. jejuni*-associated epithelial state characterized by cell-cycle checkpoint, redox, and metabolic remodeling and provide testable hypotheses for functional validation.

### Cluster composition across biological replicates

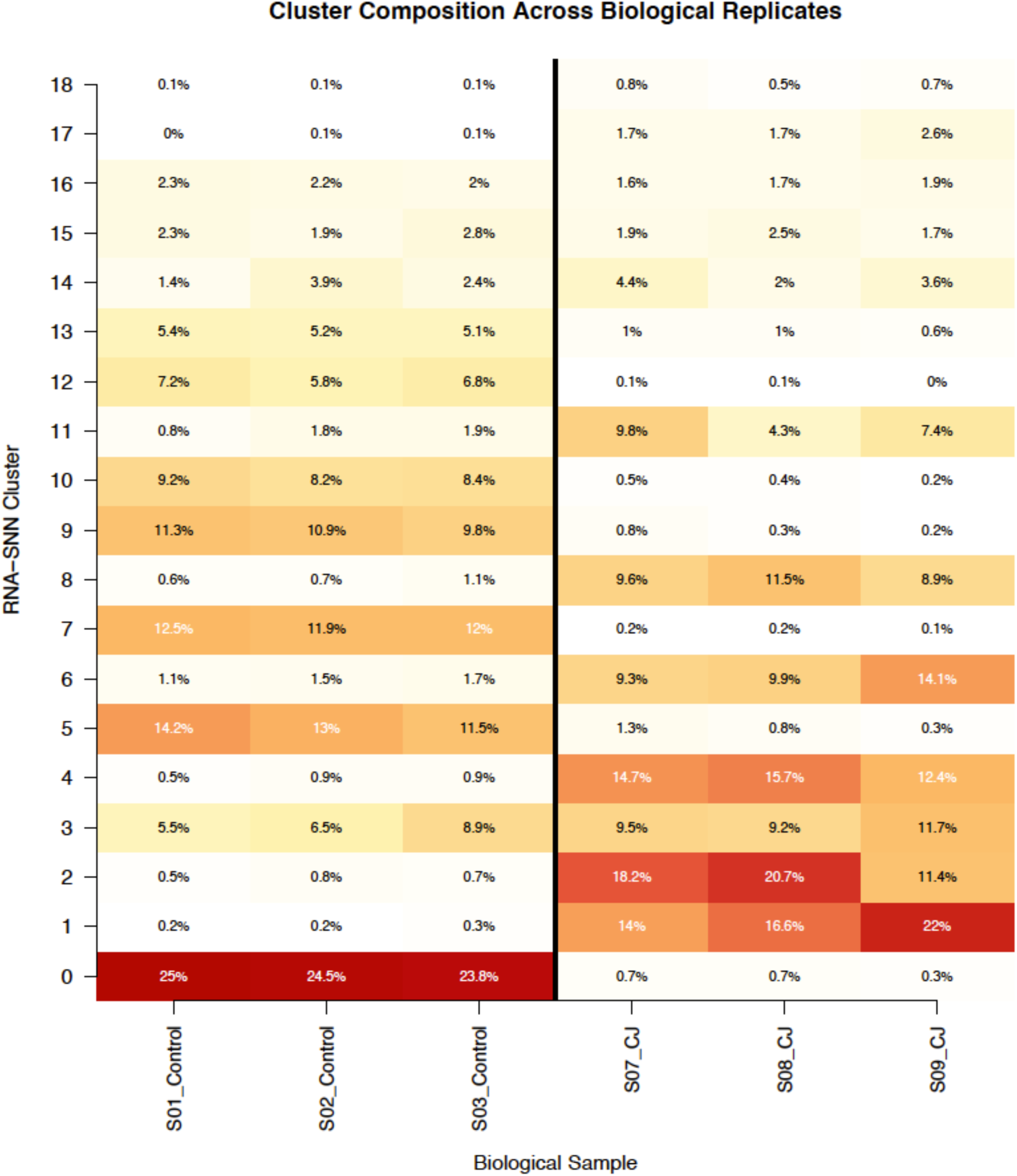

Cluster composition across biological replicates. Percent composition of the 19 RNA-SNN clusters in three Control (S01-S03) and three C. jejuni (S07-S09) samples. Control-associated clusters are reproducibly enriched in all three Controls, whereas clusters 1, 2, 4, 6, 8, and 11 expand reproducibly in infection. This pattern demonstrates that the major cluster redistribution is condition-associated rather than driven by a single sample.

### Figure: Cell-cycle and mitotic checkpoint gene violins

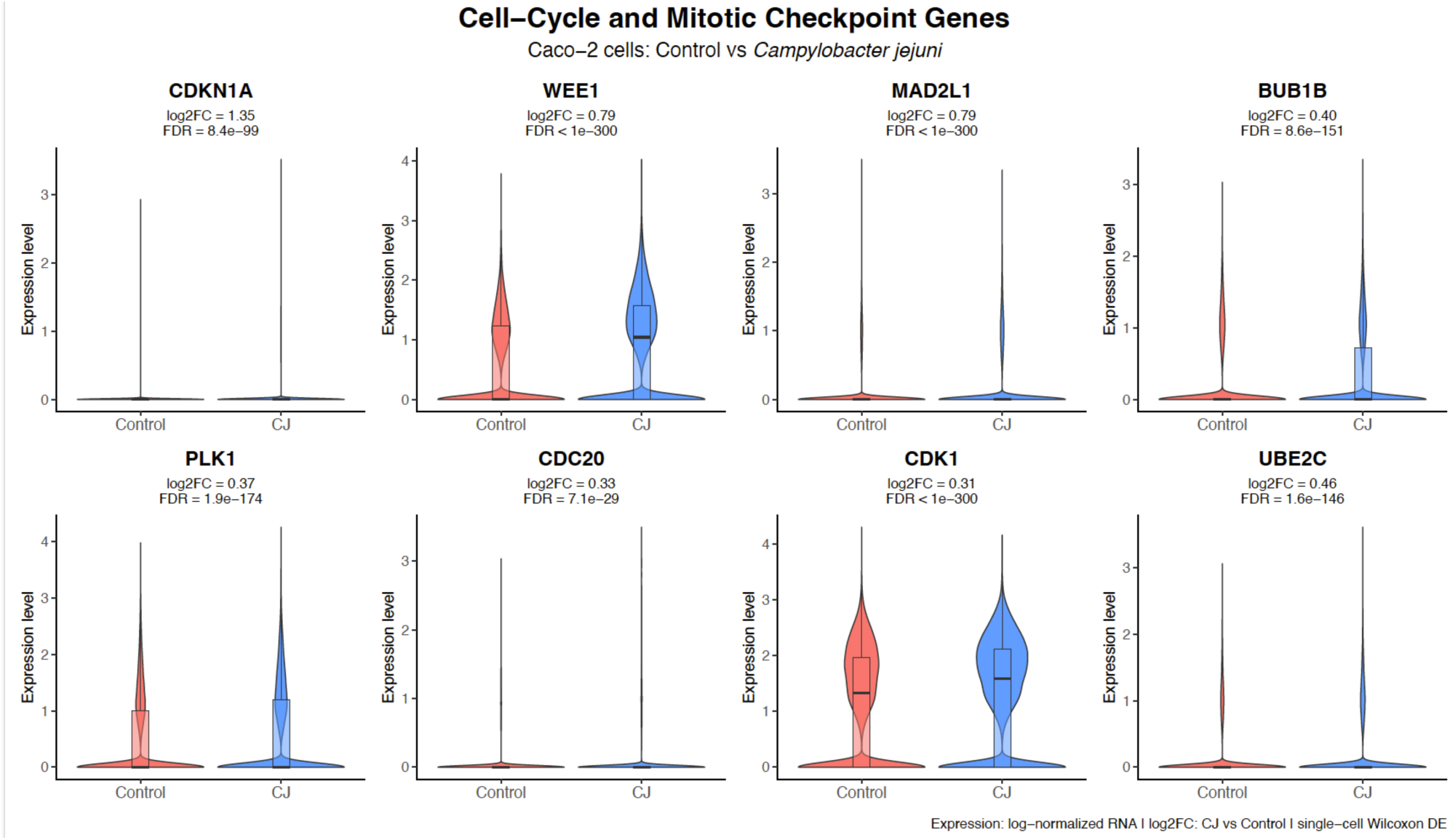

Selected checkpoint and mitotic genes in Control and C. jejuni cells. Violin plots show log-normalized expression of CDKN1A, WEE1, MAD2L1, BUB1B, PLK1, CDC20, CDK1, and UBE2C. Reported log2FC values are C. jejuni versus Control from whole-dataset Wilcoxon DE; FDR values are BH-adjusted. Several genes are zero-inflated, so differences can reflect both expression magnitude and the fraction of expressing cells.

### Figure: AUCell metabolic pathway heatmap

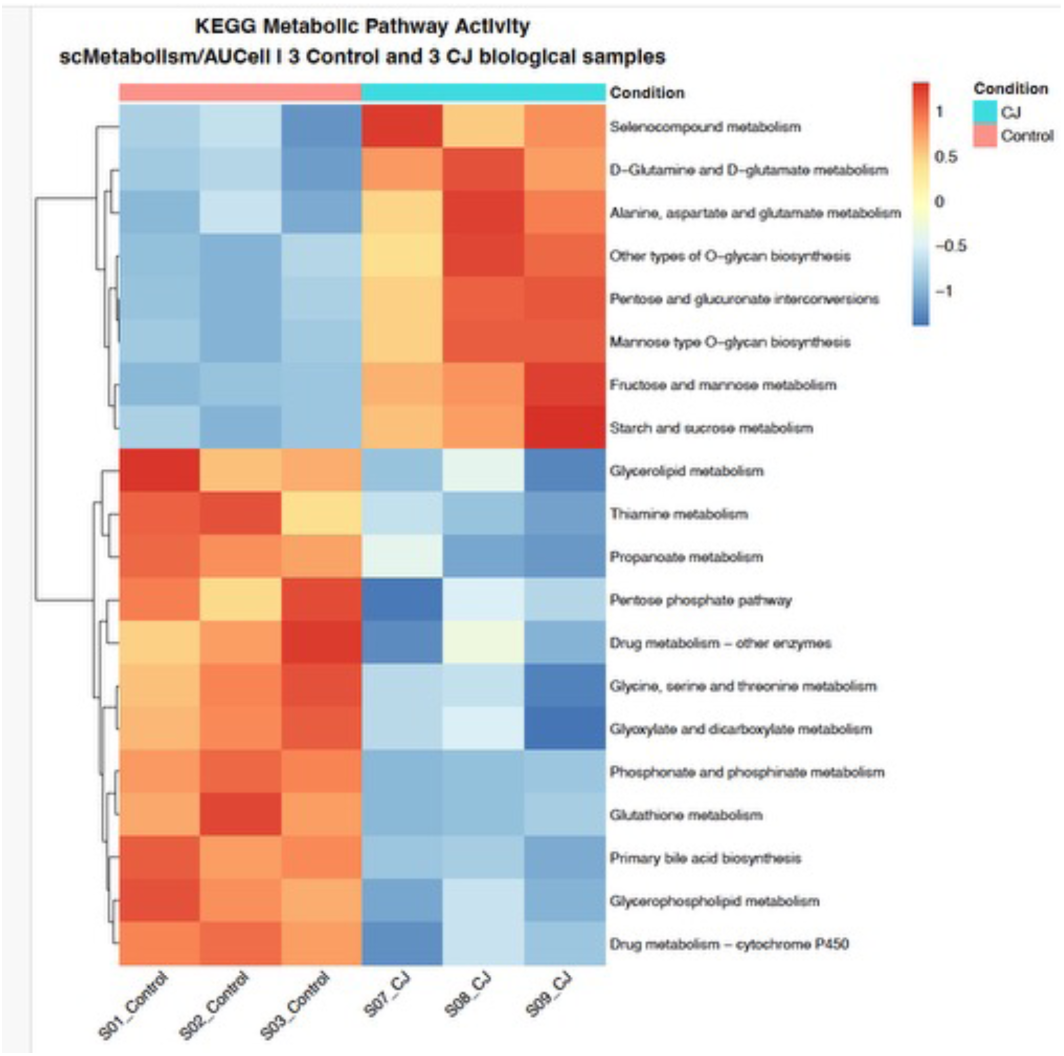

Bidirectional metabolic transcriptional remodeling. scMetabolism KEGG gene sets were scored with AUCell in a balanced 10,000-cell subset and averaged within each biological sample. Twenty of 85 pathways met FDR <0.05 in three-Control versus three-C. jejuni sample-level comparisons. Twelve were lower and eight higher in infection. Pathway-wise Z scores demonstrate strong replicate consistency. AUCell scores reflect transcriptional pathway activity, not metabolic flux.

## Notes

### Competing Interest Statement

The authors have declared no competing interest.

